# A TNF-IL-1 circuit controls *Yersinia* within intestinal granulomas

**DOI:** 10.1101/2023.04.21.537749

**Authors:** Rina Matsuda, Sorobetea Daniel, Jenna Zhang, Stefan T. Peterson, James P. Grayczyk, Beatrice Herrmann, Winslow Yost, Rosemary O’Neill, Andrea C. Bohrer, Matthew Lanza, Charles-Antoine Assenmacher, Katrin D. Mayer-Barber, Sunny Shin, Igor E. Brodsky

**Author notes:** correspondence, ORCID 0000-0001-7970-872X, ORCID 0000-0001-5214-9577. These authors contributed equally (Rina Matsuda, Sorobetea Daniel, Jenna Zhang). Current address: Oncology Discovery, AbbVie Inc., North Chicago, IL. Current address: Penn State Health Milton S Hershey Medical Center, Penn State College of Medicine, Hershey, PA 17033 USA.

## Abstract

**Summary:** Monocytes restrict *Yersinia* infection within intestinal granulomas. Here, we report that monocyte-intrinsic TNF signaling drives production of IL-1 that signals to non-hematopoietic cells to control intestinal *Yersinia* infection within granulomas.

Tumor necrosis factor (TNF) is a pleiotropic inflammatory cytokine that mediates antimicrobial defense and granuloma formation in response to infection by numerous pathogens. *Yersinia pseudotuberculosis* colonizes the intestinal mucosa and induces recruitment of neutrophils and inflammatory monocytes into organized immune structures termed pyogranulomas that control the bacterial infection. Inflammatory monocytes are essential for control and clearance of *Yersinia* within intestinal pyogranulomas, but how monocytes mediate *Yersinia* restriction is poorly understood. Here, we demonstrate that TNF signaling in monocytes is required for bacterial containment following enteric *Yersinia* infection. We further show that monocyte-intrinsic TNFR1 signaling drives production of monocyte-derived interleukin-1 (IL-1), which signals through IL-1 receptor on non-hematopoietic cells to enable pyogranuloma-mediated control of *Yersinia* infection. Altogether, our work reveals a monocyte-intrinsic TNF-IL-1 collaborative circuit as a crucial driver of intestinal granuloma function, and defines the cellular target of TNF signaling that restricts intestinal *Yersinia* infection.

## Introduction

Granulomas form in response to a wide variety of infections, acting as barriers to pathogen dissemination^1, 2^. Although generally considered protective, granulomas can also provide a replicative niche from which pathogens can spread, such as in immune-compromised patients that experience reactivation of latent *Mycobacterium tuberculosis*^3, 4^. Moreover, pathogens within granulomas often persist in an antibiotic-resistant state and can pose a significant therapeutic challenge^5^ Granulomas thus represent a localized niche within which pathogens persist and remain resistant to host immune clearance. Understanding how pathogens are controlled within granulomas remains an important question that could enable development of immunomodulatory treatments against infectious agents that persist within this niche.

Tumor necrosis factor (TNF) is a pleiotropic inflammatory cytokine associated with protection during granulomatous disease, notably tuberculosis^6–13^. While the role of TNF in maintaining intact granulomas is well-appreciated, its precise cellular targets and mechanisms of action remain elusive due to broad expression of its main receptor, TNFR1, and its pleiotropic downstream signaling functions, including induction of cell-extrinsic apoptosis, promoting cell survival, and mediating expression of pro-inflammatory gene programs^14–16^. TNF plays a critical role clinically in protection against infection by intracellular pathogens, as the extensive clinical use of anti-TNF blockade in the setting of auto-inflammatory disease is associated with increased risk of severe infection^16, 17^.

The enteropathogenic *Yersiniae*, which also include *Y. pseudotuberculosis* (*Yp*) and *Y. enterocolitica*, colonize the intestinal mucosa and lymphoid tissues of both mice and humans, triggering formation of pyogranulomas (PG) that are composed of extracellular bacterial colonies in close association with neutrophils, bordered in turn by monocytes and macrophages^18–25^. We recently demonstrated that PG containing viable bacteria, inflammatory monocytes, and neutrophils form along the length of the gastrointestinal tract early following oral *Yp* infection^25^. PG form in response to the activity of Yersinia Outer Proteins (Yops), which are injected into host cells through the *Yersinia* type III secretion system and block essential antimicrobial functions^25–27^. We further demonstrated that inflammatory monocytes were critical for maintenance of PG architecture and enabling neutrophils to overcome the activity of *Yp* virulence factors that block host phagocytosis and oxidative burst^25^. However, the mechanisms by which inflammatory monocytes mediate anti-*Yersinia* host defense are unclear.

Here, we demonstrate that monocytes serve as an essential cellular source of TNF, which is required for host protection against *Yersinia*^28–30^. We find that signaling through both TNF and IL-1 receptor are required to maintain PG control of *Yp.* Intriguingly, monocyte-intrinsic TNF production and receptor signaling were required for PG monocytes to produce IL-1, an inflammatory cytokine involved in control of other microbial infections. IL-1 in turn signals to IL-1 receptor on non-hematopoietic cells to enable control of intestinal *Yp* infection. Altogether, our study demonstrates that a monocyte-driven TNF-IL-1 signaling circuit mediates the control of *Yp* infection within systemic and intestinal sites and demonstrates that TNF and IL-1 collaborate via a feed-forward loop to promote host defense against microbial infection.

## Results

### TNFR1 is required for organized pyogranuloma formation and restriction of Yersinia

We recently identified the formation of pyogranulomas (PG) in the murine intestinal mucosa during acute *Yersinia pseudotuberculosis (Yp)* infection, wherein inflammatory monocytes were required for neutrophil activation, maintenance of PG architecture, and bacterial clearance^25^. Nonetheless, the monocyte-derived signals required for the function and maintenance of these intestinal PG are unknown. Tumor necrosis factor (TNF) is critical for granuloma maintenance and bacterial control in the lung during tuberculosis infection^6–13^, and we previously found that TNF signaling is necessary for the control of bacterial burdens following oral *Yersinia* infection^30^. Notably, while *Tnfr1^-/-^* mice formed similar numbers of macroscopic intestinal lesions as wild-type (WT) mice (Fig. S1A), histopathologic analyses revealed that intestinal lesions in *Tnfr1^-/-^*mice displayed a disorganized appearance and contained a central area of tissue necrosis that was strikingly similar to lesions that we recently described in monocyte-deficient mice^25^, (Fig. 1A). In contrast to WT PG, which had robust immune cell aggregation and a small central *Yp* microcolony, *Tnfr1^-/-^* intestinal lesions contained limited immune cell infiltrate and an expanded *Yp* colony (Fig. 1A, B). In line with these histopathological findings, bacterial burdens in pyogranuloma-containing (PG+) intestinal punch biopsies and adjacent non-pyogranuloma (PG-) biopsies were elevated in *Tnfr1^-/-^* mice (Fig. 1C). Furthermore, *Tnfr1*^-/-^ PG contained fewer monocytes, macrophages, and neutrophils, as determined by flow cytometry (Fig. 1D, S1B). Surface expression of the integrin CD11b, a marker of neutrophil activation^31–33^, was significantly reduced in PG of *Tnfr1^-/-^* mice compared to WT controls (Fig. 1E), suggesting a defect in neutrophil activation in the absence of TNFR1 signaling, consistent with our recent findings of reduced neutrophil activation within PG in the absence of monocytes^25^. Additionally, *Tnfr1^-/-^*mice exhibited elevated bacterial burdens in the spleen and liver (Fig. 1F), consistent with our previous findings^30^. Notably, *Tnfr1^-/-^* mice succumbed to infection around day 8, while most WT mice survived (Fig. 1G). Overall, these data suggest that TNFR1 signaling is necessary to mediate functional intestinal PG formation and control of *Yp*.

**Figure 1.**
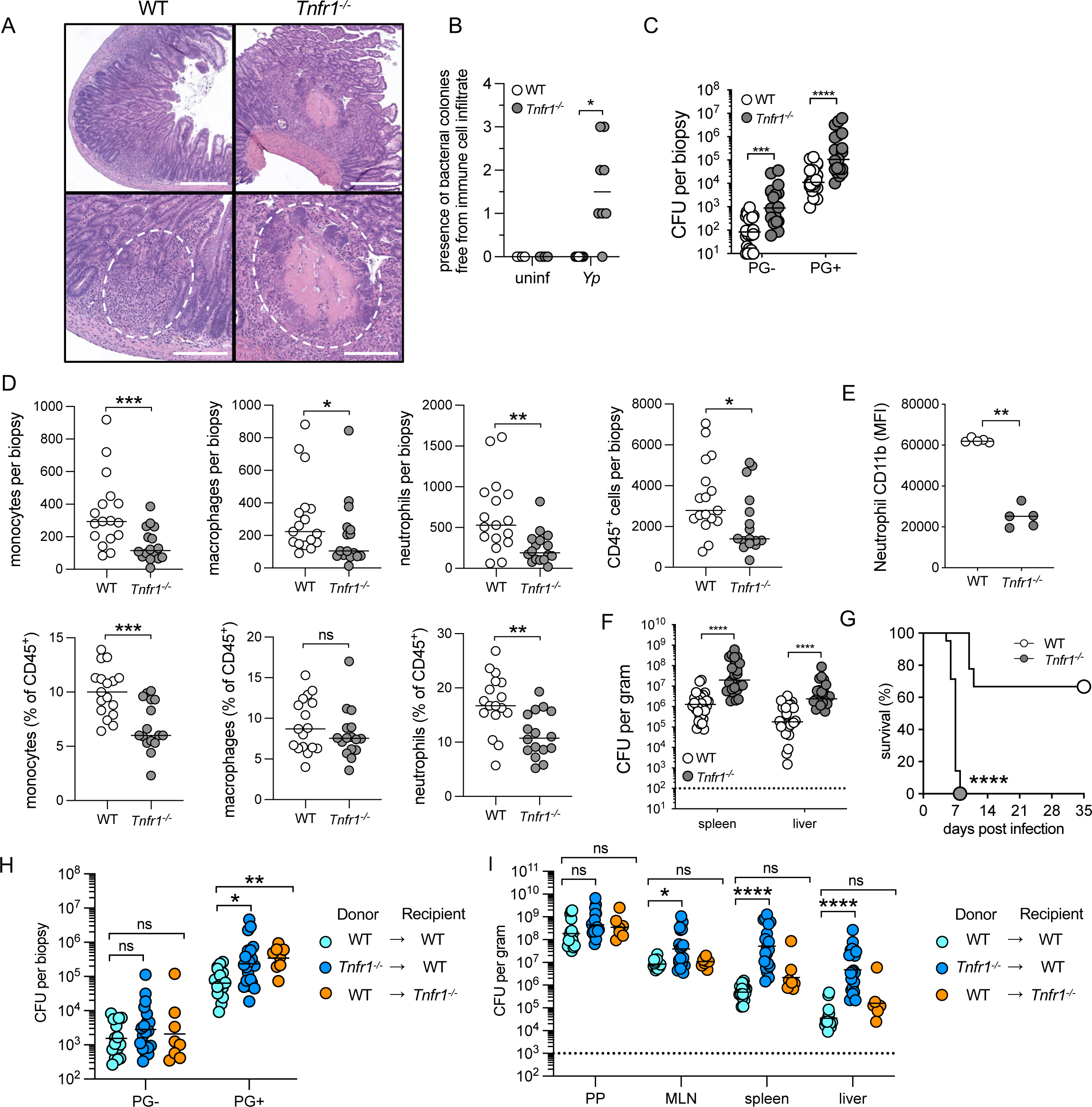
TNFR1 is required for organized pyogranuloma formation and restriction of *Yersinia* in intestine and periphery. (A) H&E-stained paraffin-embedded longitudinal small intestinal sections from *Yp*-infected mice at day 5 post-infection. Dashed line highlights pyogranuloma (left) or necrosuppurative lesion (right). Images representative of two independent experiments. Scale bars = 500 μm (top) and 200 μm (bottom). (B) Histopathological scores of small intestinal tissue from uninfected or *Yp*-infected mice at day 5 post-infection. Each mouse was scored between 0-4 (healthy-severe) for indicated sign of pathology. Each circle represents one mouse. Lines represent median. Pooled data from two independent experiments. (C) Bacterial burdens in small intestinal PG-and PG+ tissue isolated day 5 post-infection. Each circle represents the mean CFU of 3-5 pooled punch biopsies from one mouse. Lines represent geometric mean. Pooled data from three independent experiments. (D) Total numbers and frequencies of CD45^+^ cells, monocytes, macrophages, and neutrophils in uninfected, PG-, and PG+ small intestinal tissue isolated 5 days post-infection. Each circle represents the mean of 3-10 pooled punch biopsies from one mouse. Lines represent median. Pooled data from three independent experiments. (E) Mean fluorescence intensity (MFI) of CD11b expression on neutrophils in PG+ tissue at day 5 post-infection. Each circle represents the mean of 3-10 pooled punch biopsies from one mouse. Lines represent median. Data representative of three independent experiments. (F) Bacterial burdens in indicated organs at day 5 post-infection. Each circle represents one mouse. Lines represent geometric mean. Pooled data from four independent experiments. (G) Survival of infected WT (n=9) and *Tnfr1^-/-^* (n=21) mice. Pooled data from two independent experiments. (H) Bacterial burdens in small intestinal PG- and PG+ tissue at day 5 post-infection of indicated chimeric mice. Each circle represents the mean *Yp*-CFU of 3-5 pooled punch biopsies from one mouse. Lines represent geometric mean. Pooled data from two independent experiments. (I) Bacterial burdens in indicated organs at day 5 post-infection of indicated chimeric mice. Each circle represents one mouse. Lines represent geometric mean. Pooled data from two independent experiments. Statistical analysis by Mann-Whitney U test (B, C, D, E, F), Mantel-Cox test (G), and Kruskal-Wallis test with Dunn’s multiple comparisons correction (H, I) *p<0.05, **p<0.01, ***p<0.001, ****p<0.0001, ns = not significant.

TNFR1 signaling can enhance the ability of hematopoietic (immune) or non-hematopoietic (stromal) cells to control pathogens^10, 11, 13, 30, 34^. To test which compartment requires TNFR1 signaling to control *Yp*, we generated bone marrow chimeras in which TNFR1 expression was ablated on the immune or stromal compartment (Fig. S1C). Mice lacking TNFR1 in either the immune or stromal compartment had elevated bacterial burdens within PG compared to WT control chimeras, indicating that TNFR1 signaling is required non-redundantly in both hematopoietic and non-hematopoietic cells to mediate control of intestinal *Yp* (Fig. 1H). In contrast, mice lacking TNFR1 in immune cells had elevated bacterial burdens in the systemic tissues, while mice lacking TNFR1 in stromal cells had similar bacterial burdens in systemic tissues as WT controls (Fig. 1I). Taken together, these results demonstrate that TNFR1 signaling in both hematopoietic and non-hematopoietic cells contribute to bacterial control in the intestine, while TNFR1 signaling specifically in immune cells is required for bacterial control in the systemic tissues during acute *Yp* infection.

### Autocrine TNF signaling in monocytes is required for control of Yersinia

TNF receptor expression is widespread on hematopoietic cells, raising the question of which specific cells are the necessary targets of TNF signaling for control of *Yp* infection. We previously demonstrated that CCR2-deficient mice lacking circulating monocytes fail to form functional intestinal PG, are unable to control *Yp* burdens, and succumb to infection^25^. Given the similar outcomes of infection and histopathological appearance of PG in TNFR1- and CCR2-deficient mice, we sought to test the hypothesis that TNF is either produced or detected by monocytes. To do this, we generated mice in which TNFR1 was specifically deleted on inflammatory monocytes by means of mixed BM chimeras where irradiated wild-type recipient mice were reconstituted with a 1:1 ratio of *Ccr2^gfp/gfp^*:*Tnfr1^-/-^* or *Ccr2^gfp/gfp^*:*WT* control BM cells (Fig. 2A). Because circulating monocytes in these chimeric mice are derived from the *Tnfr1^-/-^*or WT BM cells, respectively, this approach generates mice in which circulating CCR2^+^ monocytes lacked or expressed TNFR1, respectively, with other hematopoietic cell types being comprised of a 1:1 mixture of these genotypes (Fig. S2A). Intriguingly, mice lacking TNFR1 specifically on CCR2^+^ monocytes formed lesions with expanded bacterial colonies and failed to control *Yp* infection, largely recapitulating the phenotype of mice lacking CCR2 in the hematopoietic system altogether (Fig. 2B-D). Importantly, this defect in bacterial control was not due to a lack of TNFR1 expression on 50% of other immune cells, as mixed chimeras from *Tnfr1^-/-^:WT* mice still had significantly lower bacterial burdens relative to *Tnfr1^-/-^:Ccr2^gfp/gfp^* mice, notably in systemic tissues (Fig. S2B, C). Mice lacking TNFR1 expression on all hematopoietic cells had significantly higher burdens than mice lacking TNFR1 on monocytes alone (Fig. S2B). Altogether, these data suggest that TNFR1 signaling in monocytes is essential for their protective role against *Yp* infection, and that TNFR1 has additional important roles in other cell types beyond monocytes.

**Figure 2.**
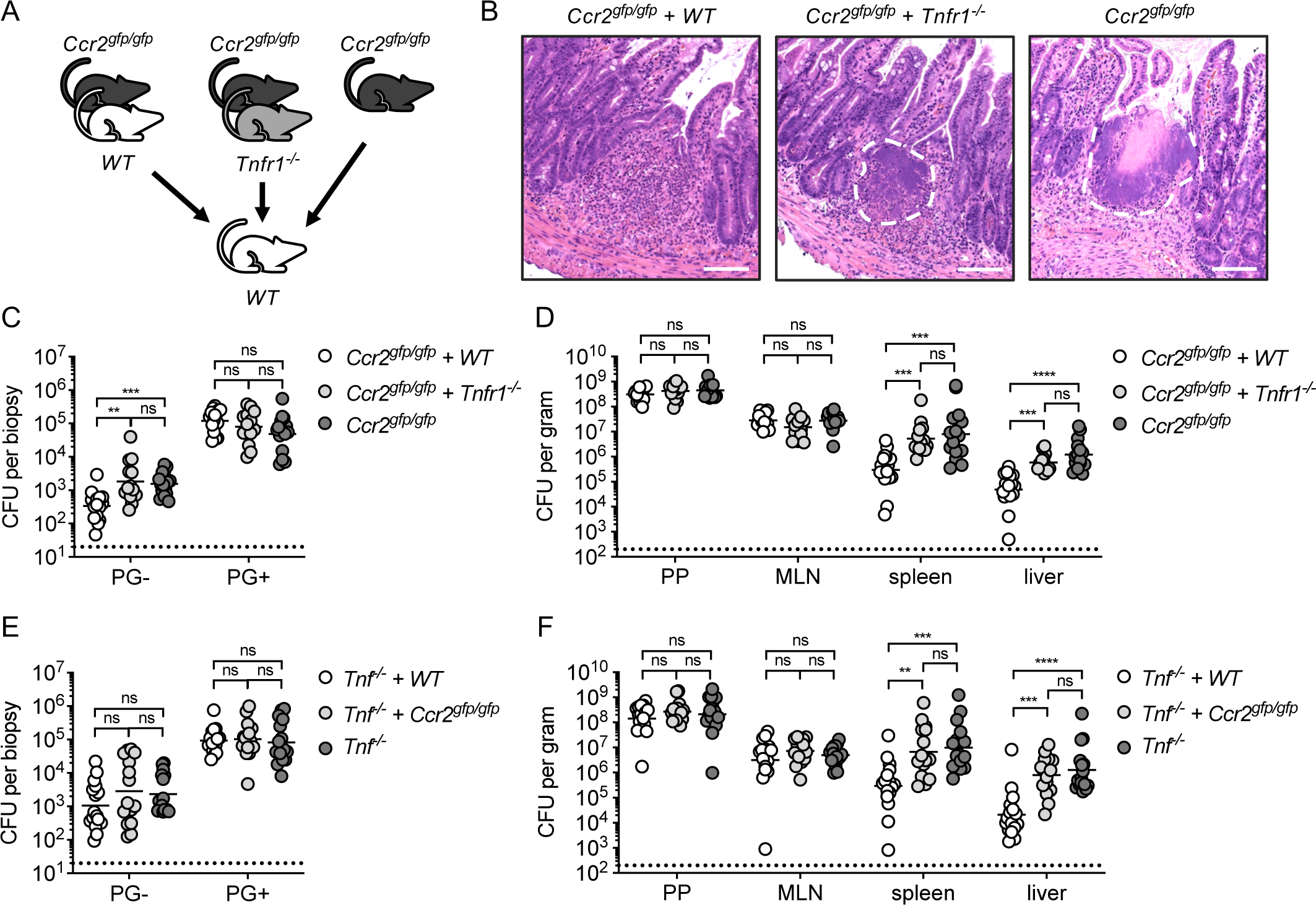
Autocrine TNF signaling in monocytes is required for control of *Yersinia*. (A) Schematic of mixed bone marrow chimeras. (B) H&E-stained paraffin-embedded transverse small-intestinal sections from chimeric WT mice reconstituted with *Ccr2^gfp/gfp^* + *WT* (left), *Ccr2^gfp/gfp^*+ *Tnfr1^-/-^* (middle), or *Ccr2^gfp/gfp^* (right) bone marrow, at day 5 post-infection. Dotted lines highlight lesions. Scale bars = 100 μm. Images representative of two independent experiments. (C) Bacterial burdens in small intestinal PG- and PG+ tissue of chimeric WT mice reconstituted with either *Ccr2^gfp/gfp^* + *WT* (white), *Ccr2^gfp/gfp^*+ *Tnfr1^-/-^* (light gray), or *Ccr2^gfp/gfp^* (dark gray) at day 5 post *Yp*-infection. Each symbol represents one mouse. Lines represent geometric mean. Pooled data from two independent experiments. (D) Bacterial burdens in indicated organs at day 5 post-infection. Each circle represents one mouse. Lines represent geometric mean. Pooled data from two independent experiments. (E) Bacterial burdens in small intestinal PG- and PG+ tissue of chimeric WT mice reconstituted with either *Tnf^-/-^* + *WT* (white), *Tnf^-/-^ + Ccr2^gfp/gfp^* (light gray), or *Tnf^-/-^* (dark gray) at day 5 post *Yp*-infection. Each symbol represents one mouse. Lines represent geometric mean. Pooled data from three independent experiments. (F) Bacterial burdens in indicated organs at day 5 post-infection. Each circle represents one mouse. Lines represent geometric mean. Pooled data from three independent experiments. Statistical analysis by Kruskal-Wallis test with Dunn’s multiple comparisons correction. *p<0.05, **p<0.01, ***p<0.001, ****p<0.0001, ns = not significant.

Multiple immune cell types produce TNF in response to inflammatory signals, including monocytes, which we previously observed to be a major source of TNF during *Yp* infection^35^. Thus, we considered that monocytes might be an important source as well as recipient of the TNF signal to enable control of *Yp* infection. We therefore reconstituted irradiated wild-type recipient mice with a 1:1 ratio of *Ccr2^gfp/gfp^*:*Tnf^-/-^*or *Tnf^-/-^:WT* control BM cells, in order to generate cohorts of mice in which circulating CCR2^+^ monocytes lacked or retained the ability to produce TNF, respectively, with other hematopoietic cell types being comprised of a 1:1 mixture. Strikingly, mice lacking TNF specifically in monocytes failed to control *Yp* infection in the spleen and liver, with equal burdens to those completely lacking TNF production in all hematopoietic cells (Fig. 2E, F). Altogether, our findings demonstrate that autocrine TNF signaling in monocytes is required to control enteric *Yp* infection.

### TNFR1 signaling in monocytes controls Yp infection independently of RIPK1 kinase-induced cell death

TNFR1 can mediate inflammatory gene expression or promote cell-extrinsic apoptosis in response to infection by pathogens, including *Yersinia*^36–41^. *Yp*-induced cell death is triggered by YopJ*-*induced blockade of IKK signaling and involves contributions from both TLR4/TRIF and TNFR1 signaling through the adapter kinase RIPK1^35, 42–47^. We previously demonstrated that mice specifically lacking RIPK1 kinase activity (*Ripk1^K45A^*) in hematopoietic cells fail to form intact MLN PG and rapidly succumb to *Yp* infection^35^.

Furthermore, activation of gasdermin D and gasdermin E in macrophages and neutrophils, respectively, downstream of RIPK1 kinase activity promotes control of *Yp* infection^48^. *Ripk1^K45A^* mice formed necrotic intestinal lesions and were deficient in restricting *Yp* burdens, consistent with prior findings (Fig. 3A-C). These data provoked the hypothesis that monocyte-intrinsic TNFR1 signaling promotes anti-*Yersinia* host defense through activation of RIPK1-induced monocyte cell death. To directly test this, we generated mixed BM chimeras in which irradiated WT recipient mice were reconstituted with a 1:1 ratio of *Ccr2^gfp/gfp^*:*Ripk1^K45A^*or *Ccr2^gfp/gfp^*:*WT* control BM cells. Following reconstitution, mice contained circulating CCR2^+^ monocytes that either lacked or expressed RIPK1 kinase activity, respectively, with all other hematopoietic cells being equally reconstituted by both donor bone marrow progenitors (Fig. S3A). Surprisingly, in contrast to hematopoietic loss of RIPK1 kinase activity, monocyte-specific ablation of RIPK1 kinase activity had no effect on the ability of mice to form intact intestinal PG or control enteric *Yp* infection (Fig. 3D-F), indicating that RIPK1 kinase activity is dispensable in monocytes to control *Yp* infection downstream of TNF signaling. Our previous findings demonstrated that the acute susceptibility of *Ripk1^K45A^* mice is reversed in the setting of infection with YopJ-deficient bacteria, illustrating that RIPK1 kinase-induced cell death is necessary to counteract the blockade of immune signaling by YopJ^35^. However, *Tnfr1^-/-^* mice still formed necrotic intestinal PG, were unable to control bacterial burdens in systemic tissues, and succumbed to infection by YopJ-deficient bacteria (Fig. 3G-I). Collectively, these data indicate TNFR1 signaling contributes to anti-*Yersinia* host defense via a mechanism distinct from RIPK1-induced cell death. We recently reported that intestinal PG form in response to the activities of the actin cytoskeleton-disrupting effectors YopE and YopH, and that monocytes counteract YopH-mediated blockade of innate immunity^25^. YopE and YopH both block phagocytosis and the oxidative burst through disruption of actin cytoskeleton rearrangement^49–58^. However, whether TNFR1 is required to overcome the immune blockade posed by YopE and YopH is unknown. Strikingly, TNFR1-deficient mice survived infection with *yopEH* mutant *Yp* (Fig. 3J), indicating that TNFR1 signaling counteracts the activity of YopE and YopH. However, in contrast to our previous findings with CCR2-deficient mice^25^, *Tnfr1^-/-^*mice were not able to control either single *Yp* mutant, although there was a significant delay in mortality in response to infection with *yopH* mutant bacteria (Fig. S3B). Together, these findings demonstrate that TNFR1-mediated restriction of enteric *Yp* infection is independent of RIPK1-induced cell death, and instead counteracts the anti-phagocytic and reactive oxygen-blocking activities of YopE and YopH.

**Figure 3.**
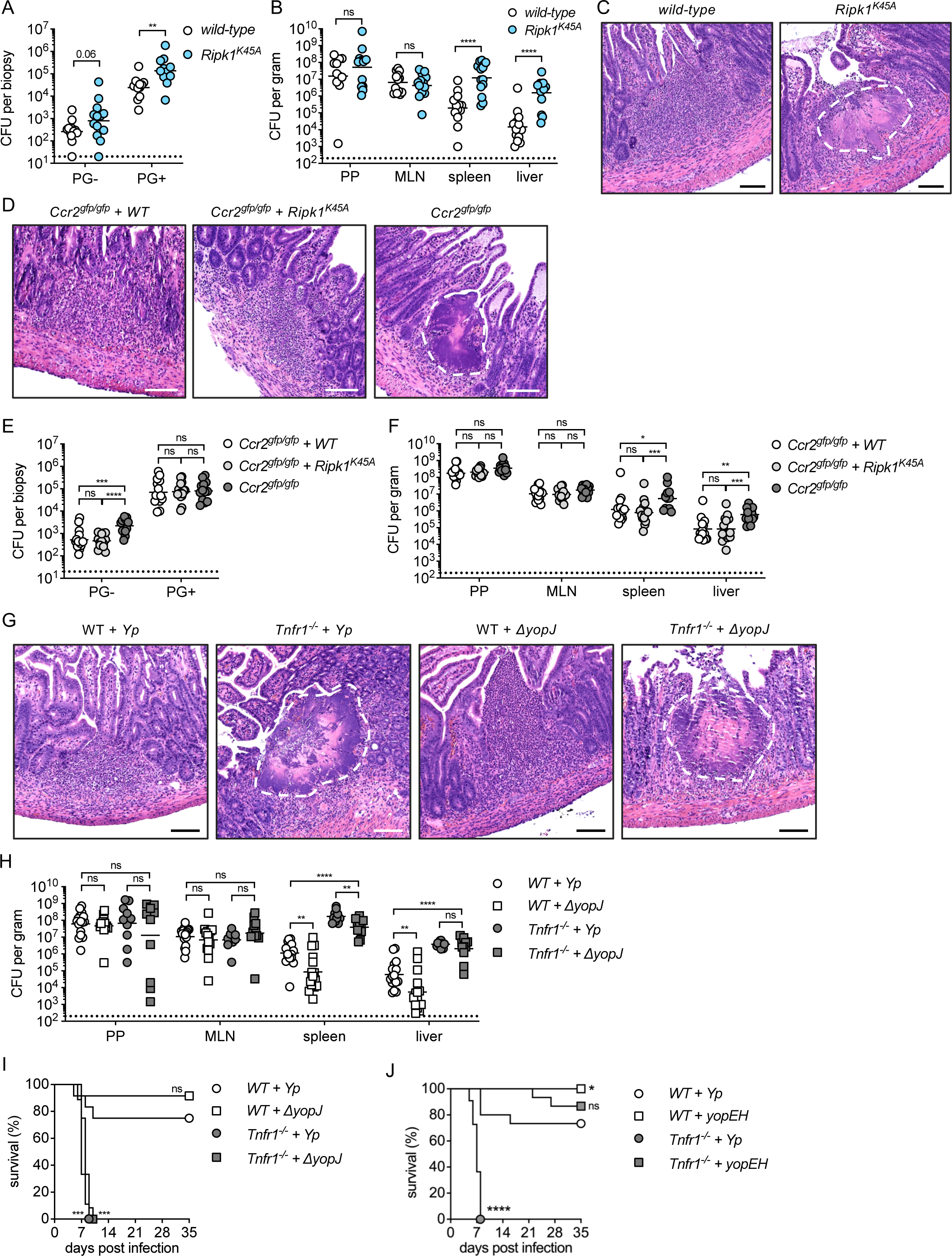
TNFR1 signaling in monocytes controls *Yp* infection independently of RIPK1 kinase-induced cell death. (A) Bacterial burdens in small intestinal PG- and PG+ tissue of WT (white) and *Ripk1^K45A^* (blue) mice at day 5 post *Yp*-infection. Each symbol represents one mouse. Lines represent geometric mean. Pooled data from two independent experiments. (B) Bacterial burdens in indicated organs at day 5 post-infection. Each circle represents one mouse. Lines represent geometric mean. Pooled data from two independent experiments. (C) H&E-stained paraffin-embedded longitudinal small intestinal sections from WT (left) and *Ripk1^K45A^*(right) mice at day 5 post *Yp*-infection with dotted line highlighting lesion. Scale bars = 100 μm. Representative images of two independent experiments. (D) H&E-stained paraffin-embedded transverse small-intestinal sections from chimeric WT mice reconstituted with either *Ccr2^gfp/gfp^* + WT (left), *Ccr2^gfp/gfp^*+ *Ripk1^K45A^* (middle), or *Ccr2^gfp/gfp^* (right) bone marrow, at day 5 post *Yp*-infection with dotted line highlighting lesion. Scale bars = 100 μm. Representative images of two independent experiments. (E) Bacterial burdens in small intestinal PG- and PG+ tissue of chimeric WT mice reconstituted with either *Ccr2^gfp/gfp^* + *WT* (white), *Ccr2^gfp/gfp^*+ *Ripk1^K45A^* (light gray), or *Ccr2^gfp/gfp^* (dark gray) at day 5 post *Yp*-infection. Each symbol represents one mouse. Lines represent geometric mean. Pooled data from two independent experiments. (F) Bacterial burdens in indicated organs at day 5 post-infection. Each circle represents one mouse. Lines represent geometric mean. Pooled data from two independent experiments. (G) H&E-stained paraffin-embedded longitudinal small intestinal sections from WT and *Tnfr1^-/-^* mice infected with either WT or Δ*yopJ Yp* at day 5 post-infection. Scale bars = 100 μm. Representative images of three independent experiments. (H) Bacterial burdens in indicated organs at day 5 post-infection. Each circle represents one mouse. Lines represent geometric mean. Pooled data from four independent experiments. (I) Survival of WT (white) and *Tnfr1^-/-^* (gray) mice infected with WT (circles) or *ΔyopJ* (squares) *Yp*. n = 9-12 mice per group. Pooled data from two independent experiments. (J) Survival of WT (white) or *Tnfr1^-/-^* (gray) mice infected with WT (circles) or *yopEH* (squares) *Yp*. n = 11-15 mice per group. Pooled data from two independent experiments. Statistical analysis by Mann-Whitney U test (A, B), Kruskal-Wallis test with Dunn’s multiple comparisons correction (E, F, H), or Mantel-Cox test (I, J). *p<0.05, **p<0.01, ***p<0.001, ****p<0.0001, ns = not significant.

### Cell-intrinsic TNFR1 signaling is required for maximal IL-1 production within intestinal pyogranulomas during Yersinia infection

Our findings indicate that while TNFR1 expression on monocytes is critical for effective intestinal PG formation and control of *Yp* infection, monocyte-intrinsic RIPK1 kinase activity is dispensable for PG formation and bacterial restriction. This suggests that RIPK1 kinase-independent mechanisms mediate monocyte-dependent control of *Yp* downstream of TNFR1 signaling. We therefore hypothesized that TNFR1 signaling in monocytes may contribute to control of *Yp* infection via promoting inflammatory cytokine production. Multiplex cytokine profiling of intestinal PG from mixed BM chimeric mice lacking monocyte-intrinsic TNFR1 expression revealed that IL-1α levels were significantly decreased, in contrast to other pro-inflammatory cytokines such as IL-6 and KC (Fig. 4A, S4A). Neither IL-1α nor IL-1β were detected in the sera of these mice, suggesting that IL-1 production in response to TNFR1 signaling is localized to intestinal tissues during *Yp* infection (Fig. S4B).

**Figure 4.**
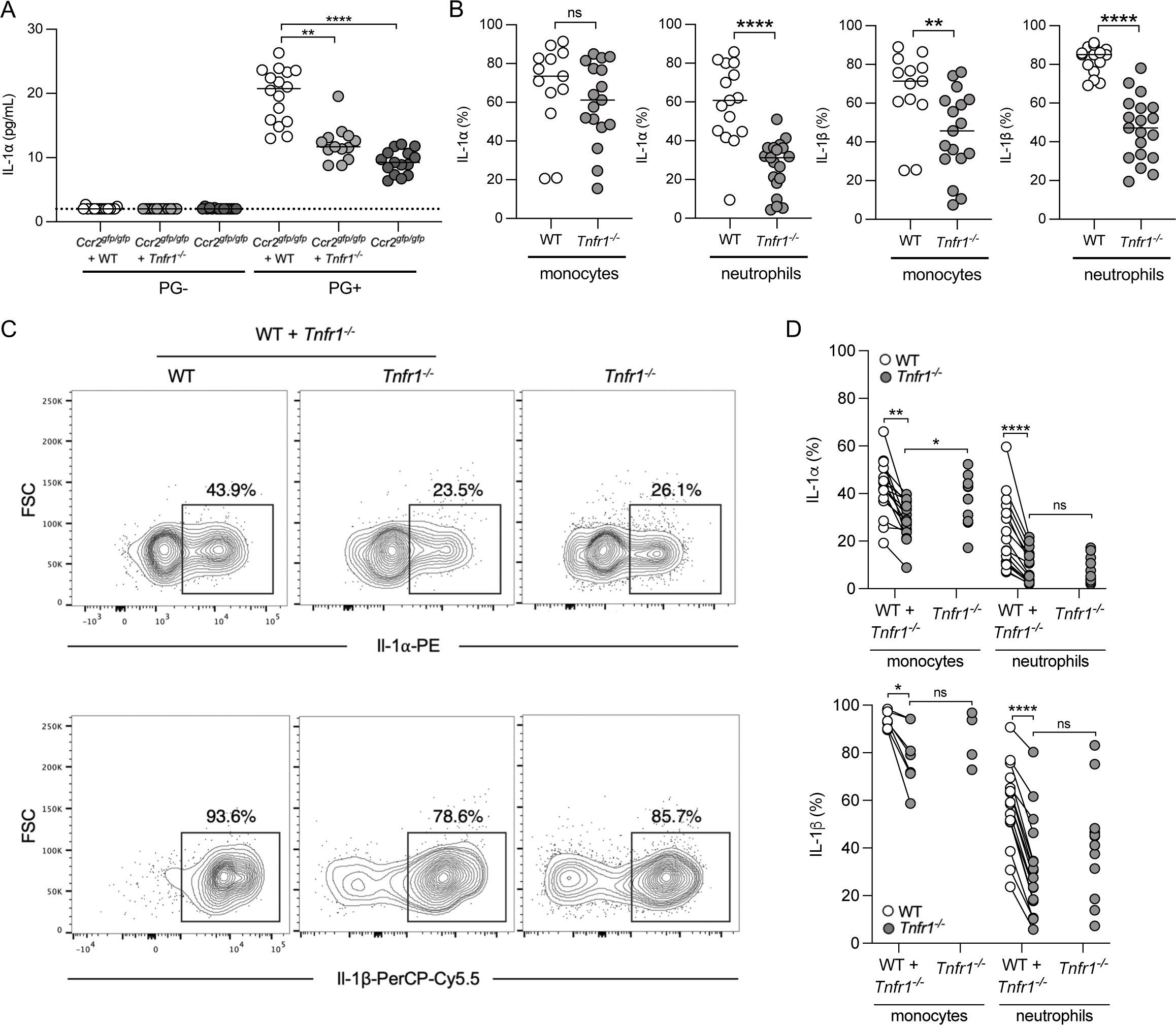
Cell-intrinsic TNFR1 signaling is required for maximal IL-1 production within intestinal pyogranulomas during *Yersinia* infection. (A) Cytokine levels were measured by cytometric bead array in tissue punch biopsy homogenates isolated 5 days post-infection from chimeric WT mice reconstituted with indicated donor cells. Lines represent median. Pooled data from two independent experiments. (B) Intracellular cytokine levels in monocytes and neutrophils isolated from small intestinal PG+ tissue 5 days post-infection. Each circle represents the mean of 3-10 pooled punch biopsies from one mouse. Lines represent median. Pooled data from three independent experiments. (C) Flow cytometry plots of intracellular IL-1 in monocytes (CD64^+^ Ly-6C^hi^) from small intestinal PG+ tissue at day 5 post-infection. Plots representative of two independent experiments. (D) Aggregate datasets from (C) for intracellular IL-1 staining in monocytes and neutrophils in small intestinal PG+ tissue at day 5 post-infection. Each circle represents the mean of 3-10 pooled punch biopsies from one mouse. Lines connect congenic cell populations within individual mice. Pooled data from two independent experiments. Statistical analysis by Kruskal-Wallis test with Dunn’s multiple comparisons correction (A), Mann-Whitney U test (B), congenic cells within mice: Wilcoxon test; across groups: Mann-Whitney U test (D). *p<0.05, **p<0.01, ***p<0.001, ****p<0.0001, ns = not significant.

Since TNFR1 expression on monocytes is required for intestinal PG formation and restriction of bacterial burdens, we hypothesized that TNFR1 signaling promotes IL-1 production by monocytes within intestinal PG. Indeed, intracellular cytokine staining demonstrated that both IL-1α and IL-1β expression were decreased in both monocytes and neutrophils in *Tnfr1^-/-^*PG, indicating that TNFR1 signaling is necessary for maximal IL-1 production in both monocytes and neutrophils within intestinal PG (Fig. 4B). We next asked whether TNFR1 signaling functions in a cell-intrinsic or-extrinsic manner to promote IL-1 cytokine production. To distinguish between these possibilities, we generated mixed bone marrow chimeras in which lethally irradiated WT recipients were reconstituted with a 1:1 mixture of WT and *Tnfr1^-/-^* bone marrow or entirely reconstituted with *Tnfr1^-/-^* bone marrow as a positive control. Importantly, there was no competitive defect in reconstitution by *Tnfr1^-/-^* cells in the mixed chimera setting, as these mice contained 1:1 ratio of WT and *Tnfr1^-/-^* immune cells within the PG and spleen (Fig. S4C). Strikingly, IL-1 production was reduced in both monocytes and neutrophils lacking TNFR1 relative to WT cells isolated from the same mice, demonstrating that cell-intrinsic TNFR1 is required for optimal production of IL-1 in monocytes and neutrophils (Fig. 4C, D, S4D). To ask if TNF signals in an autocrine fashion to upregulate its own expression, we measured intracellular TNF in these WT:*Tnfr1^-/-^* mixed chimeras. TNF levels were reduced in both monocytes and neutrophils lacking TNFR1 relative to WT cells isolated from the same intestinal PG, demonstrating that TNFR1 signals in a feedforward loop to promote TNF production in a cell-intrinsic fashion (Fig. S4E). Overall, these data show that cell-intrinsic TNFR1 signaling is necessary for maximal IL-1 production in myeloid cells within intestinal PG, raising the question of whether IL-1 production downstream of TNFR1 signaling in monocytes contributes to control of *Yp* during early intestinal infection.

### IL-1 is required for organized pyogranuloma formation and intestinal control of Yersinia

IL-1 plays a critical role in host defense by promoting immune cell recruitment and activation, cytokine production, angiogenesis, and vascular permeability^59–64^. Mice lacking IL-1 signaling are more susceptible to systemic *Yersinia* infection^48, 65, 66^. However, the role of IL-1 signaling during enteric *Yp* infection and downstream of TNF receptor signaling is unclear. IL-1β production has been proposed to promote increased intestinal permeability and barrier dysfunction^67^, suggesting multifaceted roles for IL-1 signaling within specific compartments and stages of infection. We considered the possibility that TNFR1-mediated restriction of enteric *Yp* infection and intestinal PG formation occurs in part via induction of IL-1 production from monocytes. To test the contribution of IL-1 signaling in control of enteric *Yp* infection, we infected *Il1r1*^-/-^ mice, which lack IL-1R and cannot respond to IL-1 cytokines. *Il1r1^-/-^* mice had significantly higher bacterial burdens than WT mice in the intestine, specifically in Peyer’s Patches, PG+, and PG-tissue (Fig. 5A). Notably, the intestinal lesions in *Il1r1^-/-^* mice showed extensive loss of organization and contained a central area of tissue necrosis as compared to those found in WT mice (Fig. 5B). Strikingly, the intestinal lesions in *Il1r1^-/-^* mice bore substantial resemblance to the intestinal lesions seen in *Ccr2^gfp/gfp^*mice^25^ and *Tnfr1^-/-^* mice (Fig. 5B and 1A). *Il1r1^-/-^* mice also succumbed to infection to a greater extent than WT mice (Fig. 5C). However, at day 5 post-infection, bacterial burdens in systemic organs were broadly comparable to those of WT mice (Fig. S5A). Collectively, these results suggest that consistent with PG-specific TNF-dependent IL-1 production, IL-1-mediated *Yp* restriction occurs in the intestine during early infection and that there are likely other non-IL-1-mediated mechanisms induced downstream of TNF signaling that contribute to systemic control.

**Figure 5.**
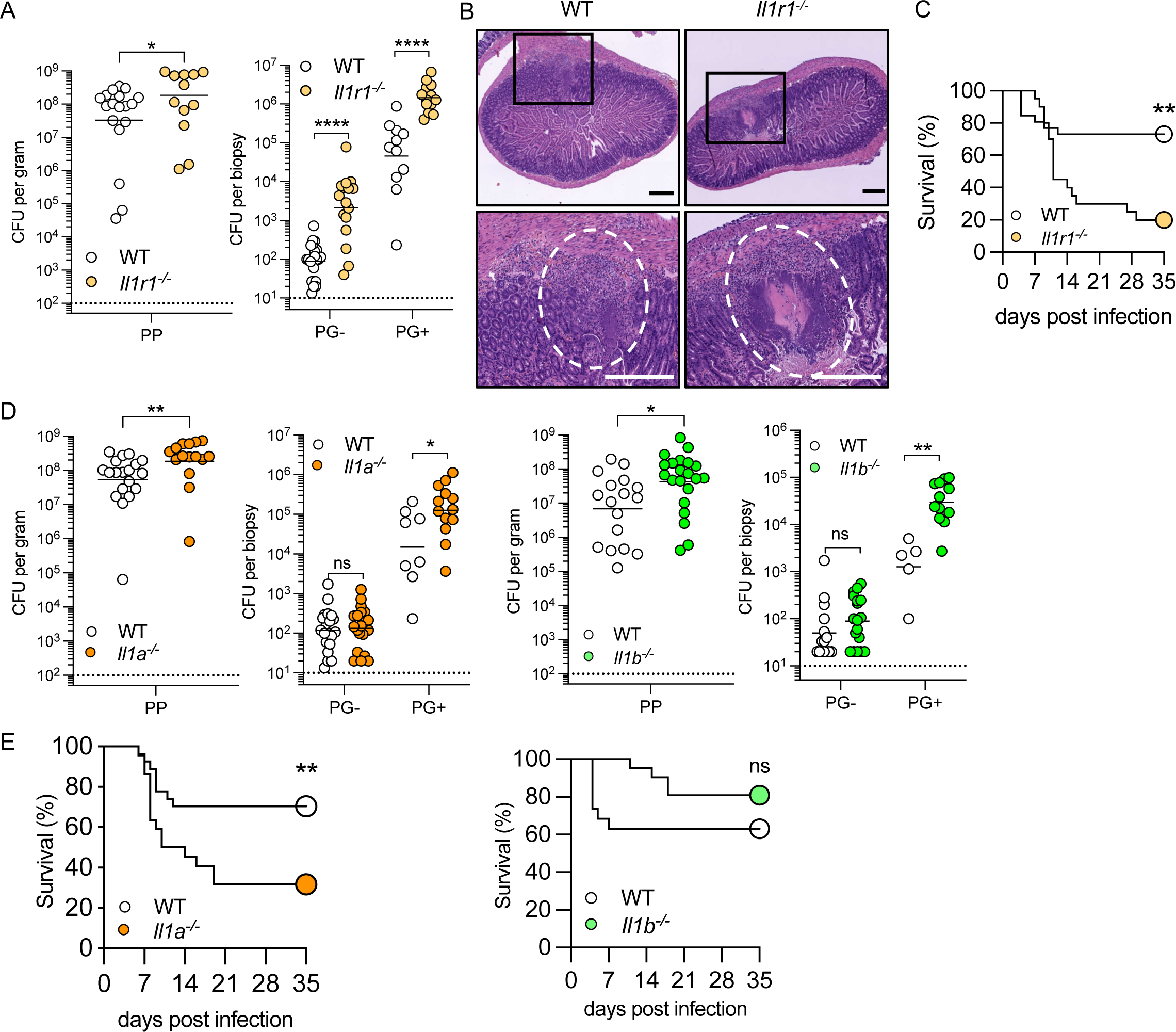
IL-1 signaling is required for organized pyogranuloma formation and intestinal control of *Yersinia*. (A) Bacterial burdens in small intestinal Peyer’s patches (PP), PG-, and PG+ tissues isolated 5 days post-infection. For PP, each circle represents pooled tissue from one mouse. For PG- and PG+, each circle represents the mean of 3-5 pooled punch biopsies from one mouse. Lines represent geometric mean. Pooled data from three independent experiments. (B) H&E-stained paraffin-embedded longitudinal small intestinal sections from *Yp*-infected mice at day 5 post-infection. Representative images of one experiment. Scale bars = 250 µm. (C) Survival of infected WT (n=26) and *Il1r1*^-/-^ (n=20) mice. Pooled data from two independent experiments (D). Bacterial burdens in small intestinal PP, PG-, and PG+ tissues at day 5 post-infection of indicated genotypes. For PP, each circle represents pooled tissue from one mouse. For PG- and PG+, each circle represents the mean of 3-5 pooled punch biopsies from one mouse. Lines represent geometric mean. Pooled data from three independent experiments. (E). Survival of infected WT (n=27, n=19), *Il1a*^-/-^ (n=22), and *Il1b*^-/-^ (n=21) mice. Pooled data from three (WT vs *Il1a*^-/-^) and two (WT vs *Il1b*^-/-^) independent experiments. Statistical analysis by Mann-Whitney U test (A, D) or Mantel-Cox test (C, E) *p<0.05, **p<0.01, ****p<0.0001, ns = not significant.

IL-1R initiates intracellular signaling cascades in response to both IL-1α and IL-1β. To test whether IL-1α and IL-1β are individually important for intestinal PG formation and *Yp* control, we infected *Il1a*^-/-^ and *Il1b*^-/-^ mice. Compared to WT mice, both *Il1a*^-/-^ and *Il1b*^-/-^ mice had elevated bacterial burdens in PP and PG+, but not in PG-tissue (Fig. 5D), in contrast to *Il1r1*^-/-^ mice which had elevated bacterial burdens in all three intestinal compartments. Like *Il1r1^-/-^* mice, *ll1a*^-/-^ and *Il1b*^-/-^ mice overall had similar bacterial burdens in systemic organs as WT mice on day 5 post-infection (Fig. S5A). Collectively, these results suggest that IL-1α and IL-1β may have overlapping roles in restricting early enteric *Yp* infection, and in the absence of one, the other may compensate. Intriguingly, *Il1a^-/-^* mice had a comparable survival defect to *Il1r1^-/-^* mice, whereas *Il1b*^-/-^ mice were similar to WT mice in survival following *Yp* infection (Fig. 5E). Collectively, these results indicate that IL-1R signaling is important for intestinal PG formation and control of enteric *Yp* infection, and may constitute a mechanism by which TNFR1 signaling controls local intestinal infection.

### Monocyte-derived IL-1 signals to non-hematopoietic cells to restrict Yersinia in intestinal pyogranulomas

Our findings demonstrate that with PG, autocrine TNF signaling in inflammatory monocytes promotes cell-intrinsic IL-1 production and subsequent IL-1R signaling promotes anti-*Yp* immune defense. However, whether monocyte-derived IL-1 is specifically required for control of intestinal *Yp* has not been tested. We therefore infected mixed BM chimeras in which irradiated wild-type recipient mice were reconstituted with a 1:1 ratio of *Il1ab*^-/-^:*Ccr2^gfp/gfp^*bone marrow cells to generate cohorts of mice specifically lacking IL-1α and IL-1β production in monocytes, along with mice reconstituted with *Il1ab*^-/-^:WT bone marrow or 100% *Il1ab^-/-^* bone marrow (Fig. S6A). Critically, mice specifically lacking IL-1α and IL-1β in monocytes had significantly elevated bacterial burdens in PG, recapitulating elevated bacteria burdens in PG of hematopoietic-deficient IL-1α and IL-1β chimeric mice and indicating that monocyte-derived IL-1 is important for restricting infection within intestinal PG (Fig. 6A). Bacterial burdens in PG-punch biopsies and systemic organs were broadly similar across chimeric mice genotypes (Fig. 6A, S6B), suggesting that IL-1 production from other cell types besides monocytes may contribute to intestinal infection restriction. Collectively, these data suggest that multiple cellular sources of IL-1 drive restriction of *Yp*. In agreement with our previous finding that TNFR1-deficient mice have a defect in IL-1 production within PG, monocyte-derived IL-1 was critical for control of bacterial burdens within PG, indicating that IL-1 production from monocytes plays a significant role in TNF-dependent control within this intestinal niche.

**Figure 6.**
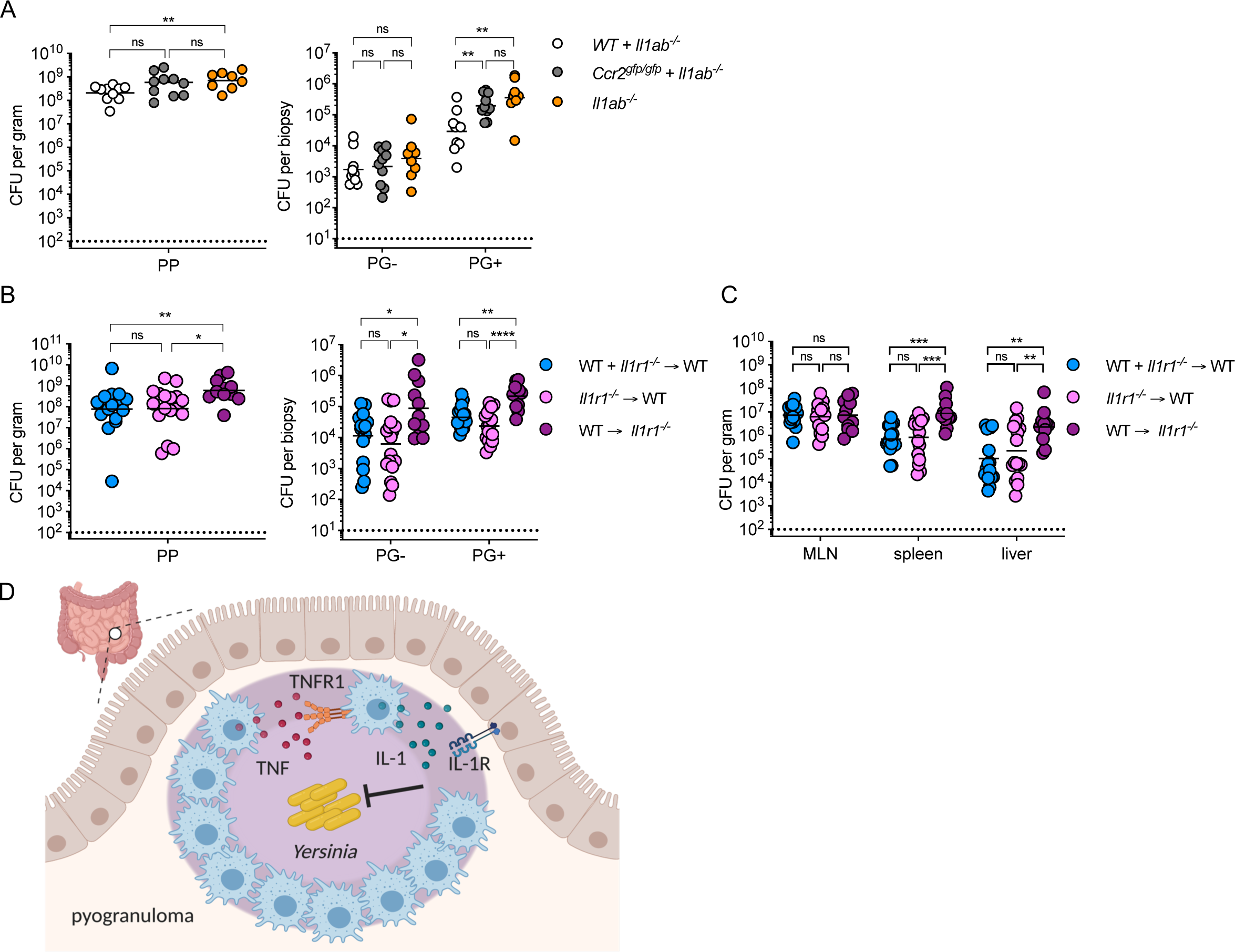
Monocyte-derived IL-1 signals to nonhematopoietic cells to restrict ***Yersinia* infection in intestinal pyogranulomas**. (**A**) Bacterial burdens in small intestinal Peyer’s patches (PP), PG-, and PG+ tissues at day 5 post-infection of indicated chimeric mice. For PP, each circle represents pooled tissue from one mouse. For PG- and PG+, each circle represents the mean of 3-5 pooled punch biopsies from one mouse. Lines represent geometric mean. Data pooled from two independent experiments. (**C**) Bacterial burdens in small intestinal Peyer’s patches (PP), PG-, and PG+ tissues isolated 5 days post-infection of indicated chimeric mice. For PP, each circle represents one mouse. For PG- and PG+, each circle represents the mean of 3-5 pooled punch biopsies from one mouse. Lines represent geometric mean. Pooled data from three independent experiments. (C) Bacterial burdens in indicated organs at day 5 post-infection of indicated chimeric mouse. Each circle represents one mouse. Lines represent geometric mean. Data pooled from three independent experiments. (D) Model of TNF-IL-1 circuit mediated by monocyte and stromal compartment to promote *Yp* restriction within intestinal pyogranulomas. All statistical analysis by Kruskal-Wallis test with Dunn’s multiple comparisons correction. *p<0.05, **p<0.01, ***p<0.001, ns = not significant.

Multiple cell types express *Il1r1* and respond to IL-1 signaling. In other infectious settings, IL-1R signaling specifically in the stromal compartment is critical for antibacterial defense^62, 64, 68–71^. We therefore considered that IL-1R signaling in stromal cells may be critical for formation and maintenance of intestinal PG as well as *Yp* restriction. To test this, we generated BM chimeric mice in which irradiated *Il1r1*^-/-^ mice were reconstituted with WT BM. Additionally, irradiated WT mice were reconstituted with either *Il1r1*^-/-^ bone marrow or a 1:1 ratio of *Il1r1*^-/-^:WT bone marrow. Notably, mice lacking IL-1R in the stromal compartment had elevated bacterial burdens in both intestinal (PP, PG+ and PG) and systemic (liver and spleen) tissues following oral *Yp* infection compared to WT control chimeras, while mice lacking IL-1R in the hematopoietic compartment had similar bacterial burdens to WT control chimeras (Fig. 6B, C). Overall, our findings demonstrate that IL-1R signaling on the stromal compartment is required to restrict *Yp* infection both in the intestinal and systemic tissues.

## Discussion

Granulomas are organized biological structures containing multiple immune cell types working in concert with stromal cells to sequester pathogens that are difficult to clear^1, 2^. *Yersinia pseudotuberculosis* induces the formation of granulomatous lesions in both the human and murine intestine^18–25^. Here, we uncover a TNF/IL-1 signaling circuit that promotes restriction of enteropathogenic *Yp* within intestinal pyogranulomas. Notably, we find that autocrine TNF signaling on inflammatory monocytes was necessary to promote cell-intrinsic IL-1 production, which signaled on the non-hematopoietic compartment to elicit control of *Yp* within intestinal pyogranulomas.

TNF has a well-established role in granuloma formation and maintenance^6–13^. Anti-TNF therapy triggers reactivation of dormant *Mycobacterium tuberculosis* infection^8, 72, 73^, and TNF promotes macrophage-dependent control of *Salmonella* replication within granulomas during chronic *Salmonella* infection^34^. TNF promotes multiple antimicrobial activities of macrophages that are critical for granuloma formation and control of tuberculosis^10, 11, 13, 74^. However, excessive TNF causes macrophage cell death that can be detrimental to bacterial control^75^. Some pathogens counteract the pro-inflammatory effects of TNF signaling within granulomas. *Salmonella* injects the T3SS effector SteE to induce anti-inflammatory M2 macrophage polarization, countering TNF-driven M1 polarization to hinder bacterial clearance^34^. In line with our previous observations that TNFR1 is required for protection against *Yp*^30^, our findings highlight a key role for TNF signaling in promoting intestinal pyogranuloma formation and function during enteropathogenic *Yp* infection.

TNFR1 signaling promotes *Yp-*induced cell death via RIPK1 activity, and we previously found that RIPK1 activity was necessary for control of *Yp* infection^35^. However, while deficiency in RIPK1 kinase activity led to disrupted intestinal PG formation during early infection, we surprisingly found that RIPK1 kinase activity was dispensable in monocyte-lineage cells for PG formation and bacterial restriction. These findings suggest that two distinct pathways are necessary for protection against enteric *Yersinia* infection:

1) a monocyte-intrinsic TNFR1 pathway that amplifies inflammatory cytokine production in monocytes, and 2) a YopJ-induced RIPK1 kinase-mediated cell death of non-monocyte cells. Notably, neutrophils undergo GSDME-dependent pyroptosis downstream of RIPK1 kinase activity, which contributes to restriction of *Yp* infection *in vivo*^48^. Future studies will elucidate whether RIPK1 kinase-dependent neutrophil cell death promotes control of *Yp*. We found that TNFR1 signaling on PG monocytes enhanced cell-intrinsic IL-1 production, consistent with our previous findings that monocyte-deficient intestinal PG have reduced IL-1 levels^25^. While TNF receptor signaling amplifies inflammasome activation, an important step in IL-1 processing^76–78^ very few studies have described TNFR-signaling-mediated IL-1 production specifically. In the context of *Legionella pneumophila* infection, IL-1 signaling induces TNF production in uninjected bystander cells in order to overcome virulence-induced host protein blockade that prevents TNF production from infected cells^61, 62^ Interestingly, *Yersinia* YopJ suppresses inflammatory cytokine expression including TNF expression^43^ Whether uninjected bystander monocytes are the critical source of TNF during enteric *Yp* infection remains unexplored. TNF has been shown to induce macrophage polarization and promotion of IL-1β expression via sterol response element binding factors^16^ How TNF receptor signaling augments IL-1 production during *Yp* infection is still unknown.

IL-1R signaling is critical for infection control during tuberculosis infection and loss of IL-1R signaling leads to expansion of pathologic lesions in the lung^79–84^. We observed that in the absence of IL-1R, mice failed to form organized intestinal pyogranulomas and had elevated intestinal *Yp* burdens, corroborating previous reports that IL-1R signaling promotes anti-*Yersinia* defense^48, 65, 66^. Intriguingly, systemic bacterial burdens were similar between WT mice and mice lacking IL-1R signaling, suggesting that there TNF signaling induces other mechanisms of bacterial restriction beyond IL-1-mediated protection. Mice lacking IL-1R signaling were more susceptible than mice lacking IL-1α or IL-1β alone, indicating that both cytokines contribute non-redundantly to bacterial restriction. While mice deficient in IL-1α exhibited elevated mortality, there was no difference in mortality in mice deficient in IL-1β. Perhaps during the early intestinal stage of infection, IL-1α and IL-1β mediate overlapping mechanisms of *Yersinia* restriction, while at later stages of infection, IL-1α is largely responsible for IL-1R-mediated control. IL-1α is a critical mediator of intestinal inflammation and inflammatory cell recruitment during *Yersinia enterocolitica* infection^85^. In contrast, IL-1β contributes to *Yersinia* restriction^48^ but also promotes intestinal barrier permeability and translocation of commensal bacteria downstream of YopJ activity^86^. Together, these observations suggest that tight regulation of intestinal IL-1 signaling is important to combat *Yersinia* infection while avoiding excessive tissue damage and loss of intestinal barrier function.

While IL-1β is released from hematopoietic cells downstream of inflammasome activation, IL-1α is more broadly expressed across cell types and can function in multiple locations, including within the nucleus, as a membrane-bound cytokine, or as an alarmin released from dying cells^87^. We found that monocyte-intrinsic IL-1α and IL-1β were necessary for control of *Yp* burdens within intestinal PG, consistent with other infectious contexts where hematopoietic-derived IL-1 drives pathogen restriction^62, 82^. However, IL-1α and IL-1β production from monocytes was dispensable for control of *Yp* burdens in PG-tissue and systemic organs, suggesting that production from other cell types contribute to IL-1-mediated infection restriction in these compartments. TNFR1 signaling also promoted neutrophil IL-1 production during enteric *Yp* infection, in line with prior studies identifying a role for GSDME-dependent IL-1β production by neutrophils in control of enteric *Yp* infection^48^. Whether neutrophils rely on a similar TNFR1-IL-1 signaling pathway to elicit control of *Yp* remains to be investigated.

Finally, IL-1R on stromal cells was critical for control of *Yp* infection. In other infectious contexts, IL-1R signaling in stromal cells is important for pathogen control, highlighting a recurring theme of IL-1R signaling cross-talk between immune and non-immune cells during infection^62, 64, 68, 88, 89^. Stromal cells are increasingly appreciated as critical components of the innate immune response. IL-1R is expressed in non-lymphoid tissues, including epithelial and endothelial cells, across various organs^90^, suggesting a conserved mechanism by which hematopoietic cytokine signaling can be amplified during infection and inflammation. The stromal cells in the intestine that respond to IL-1R signaling and the downstream anti-bacterial functions that promote restriction of *Yersinia* remain unknown. Intestinal epithelial cells respond to IL-1R signaling by upregulating antimicrobial peptide production, promoting neutrophil recruitment, and modulating intestinal permeability^59, 64, 69, 70, 89, 91, 92^. Neutrophil activation is critical within intestinal PG during *Yersinia* infection and decreased neutrophil recruitment is observed in the absence of monocytes^25^ and TNFR1 signaling. Whether IL-1R signaling on the intestinal epithelial or endothelial compartment promotes neutrophil recruitment and function during *Yp* infection remains to be determined in future studies. Altogether, our work uncovers a monocyte-intrinsic TNF/IL-1 circuit that signals to IL-1R on stromal cells to control *Yersinia* infection, providing new mechanistic insight into the cytokine networks that promote enteric granuloma formation and function.

## Methods

### Mice

C57BL/6J (CD45.2), C57BL/6.SJL (CD45.1), *Ccr2^gfp/gfp^* mice^93^ were obtained from the Jackson Laboratory. *Tnfr1^-/-^* ^94^, *Ripk1^K45A^* ^95^, *Il1r1^-/-^* ^96^, *Il1a^-/-^*^97^*, Il1b^-/-^*^97^ and *Il1a^-/-^Il1b^-/-^* ^97^ mice were previously described. All mice were bred at the University of Pennsylvania by homozygous mating and housed separately by genotype. Mice of either sex between 8-12 weeks of age were used for all experiments. All animal studies were performed in strict accordance with University of Pennsylvania Institutional Animal Care and Use Committee-approved protocols (protocol #804523).

### Bacteria

Wild-type *Yp* (clinical isolate strain 32777, serogroup O1)^98^ and isogenic YopJ-deficient mutant were provided by Dr. James Bliska (Dartmouth College) and previously described^47^. Generation of mutants lacking YopE (*ΔyopE*), enzymatic activity of YopH (YopH^R409A^), or both (denoted *yopEH*) were previously described^25^.

### Bone marrow chimeras

Wild-type B6.SJL mice (CD45.1 background) or knockout mice (*Tnfr1^-/-^* or *Il1r1^-/-^*, CD45.2 background) were lethally irradiated (1096 rads). 6 hours later, mice were injected retro-orbitally with freshly isolated bone marrow cells (5×10^6^ total cells, 2.5×10^6^ cells per donor in mixed groups) from isogenic donors of the indicated genotypes. All chimeras were provided with antibiotic-containing acidified water (40 mg trimethoprim and 200 mg sulfamethoxazole per 500 mL drinking water) for four weeks after irradiation and subsequently provided acidified water without antibiotics for a total of at least ten weeks. The reconstitution of hematopoietic cells (proportion of donor CD45^+^ cells among total CD45^+^ cells) in the blood, spleen, or intestine was analyzed by flow cytometry.

### Mouse infections

*Yp* was cultured to stationary phase at 28°C and 250 rpm shaking for 16 hours in 2×YT broth supplemented with 2 μg/ml triclosan (Millipore Sigma). Mice were fasted for 16 hours and subsequently inoculated by oral gavage with 200 μl phosphate-buffered saline (PBS) as previously^25^ All bacterial strains were administered at 2×10^8^ colony-forming units (CFU) per mouse.

### Bacterial CFU quantifications

Tissues were collected in sterile PBS, weighed, homogenized for 40 seconds with 6.35 mm ceramic spheres (MP Biomedical) using a FastPrep-24 bead beater (MP Biomedical). Samples were serially diluted tenfold in PBS, plated on LB agar supplemented with 2 μg/ml triclosan, and incubated for two days at room temperature. Dilutions of each sample were plated in triplicate and expressed as the mean CFU per gram or per biopsy.

### Cytokine quantification

Cytokines were measured in homogenized tissue supernatants using a Cytometric Bead Array (BD Biosciences) according to manufacturer’s instructions with the following modification: the amounts of capture beads, detection reagents, and sample volumes were scaled down tenfold. Data were collected on an LSRFortessa flow cytometer (BD Biosciences) and analyzed with FlowJo v10 (BD Biosciences).

### Tissue preparation and cell isolation

Blood was harvested by cardiac puncture upon euthanasia and collected in 250 U/ml Heparin solution (Millipore Sigma). Erythrocytes were lysed with Red Blood Cell Lysing Buffer (Millipore Sigma).

Spleens were homogenized through a 70 μm cell strainer (Fisher Scientific), then flushed with R10 buffer consisting of RPMI 1640 (Millipore Sigma) supplemented with 10 mM HEPES (Millipore Sigma), 10% fetal bovine serum (Omega Scientific), 1 mM sodium pyruvate (Thermo-Fisher Scientific), and 100 U/ml penicillin + 100 μg/ml streptomycin (Thermo Fisher Scientific). Erythrocytes were lysed with Red Blood Cell Lysing Buffer (Millipore Sigma).

Intestines were excised, flushed luminally with sterile PBS to remove the feces, opened longitudinally along the mesenteric side and placed luminal side down on cutting boards (Epicurean). Small intestinal tissue containing macroscopically visible pyogranulomas (PG+), adjacent non-granulomatous areas (PG-) and uninfected control tissue (uninf) were excised using a 2 mm-ø dermal punch-biopsy tool (Keyes). Biopsies within each mouse were pooled groupwise, suspended in epithelial dissociation buffer consisting of calcium and magnesium-free HBSS (Thermo Fisher Scientific) supplemented with 15 mM HEPES, 10 mg/ml bovine serum albumin (Millipore Sigma), 5 mM EDTA (Millipore Sigma), and 100 U/ml penicillin + 100 μg/ml streptomycin, then incubated for 30 minutes at 37°C under continuous agitation at 300 RPM. To isolate immune cells from the lamina propria, the tissue was enzymatically digested in R10 buffer, along with 0.5 Wünsch units/ml liberase TM (Roche), 30 μg/ml DNase I (Roche), and 5 mM CaCl_2_ for 20 min at 37°C under continuous agitation. The resulting cell suspensions were filtered through 100 μm cell strainers (Fisher Scientific) and subjected to density gradient centrifugation using Percoll (GE Healthcare). Briefly, cells were suspended in 40% Percoll and centrifuged over a 70% Percoll layer for 20 min at 600 × g with the lowest brake at room temperature. Cells collected between the layers were washed with R10 buffer for downstream analysis.

### Flow cytometry

Non-specific Fc binding was blocked for 10 minutes on ice with unconjugated anti-CD16/CD32 (93; Thermo-Fisher Scientific). Cells were subsequently labeled for 30 minutes on ice with the following antibodies and reagents: PE-conjugated rat anti-mouse Siglec-F (E50-2440; BD Biosciences), PE-TxR or PE-Cy5-conjugated rat anti-mouse CD11b (M1/70.15; Thermo Fisher Scientific), PE-Cy5.5 or PE-Cy7-conjugated rat anti-mouse CD4 (RM4-5; Thermo Fisher Scientific), BV510-conjugated rat anti-mouse CD3e (145-2C11; BioLegend), AF700 or PerCP-Cy5.5-conjugated rat anti-mouse Ly-6C (HK1.4; Thermo Fisher Scientific), BV605-conjugated Armenian hamster anti-mouse TCRβ (H57-597; BD Biosciences), BV650-conjugated rat anti-mouse I-A/I-E (M5/114.15.2; BD Biosciences), BV711-conjugated rat anti-mouse CD8α (53-6.7; BD Biosciences), BV785-conjugated rat anti-mouse Ly-6G (1A8; Thermo Fisher Scientific), PE-Cy7 or AF647-conjugated mouse anti-mouse CD64 (X54-5/7.1; BD Biosciences), AF700-conjugated mouse anti-mouse CD45.2 (104; BioLegend), PE-Cy5-conjugated mouse anti-mouse CD45.1 (A20; Thermofisher), PE-Cy5 or PE-CF594-conjugated rat anti-mouse CD45R/B220 (RA3-6B2; BD Biosciences) along with eF780 viability dye (BioLegend) diluted in PBS. Antibodies were used at 1:200 dilution and viability dye at 1:1500 dilution.

For intracellular staining, cells were incubated for 3 hours at 37°C with 5% CO_2_ in R10 buffer supplemented with 0.33 μl/ml GolgiStop (BD Biosciences) and 15 μg/ml DNase I. Surface proteins were stained as above, and cells were fixed for 20 minutes on ice with Cytofix/Cytoperm Fixation/Permeabilization solution (BD Biosciences). Intracellular cytokines were stained at 4°C overnight with FITC or PerCP-e710-conjugated rat anti-mouse IL-1β (NJTEN3; Thermo Fisher Scientific) and PE-conjugated Armenian hamster anti-mouse IL-1α (ALF-161; BioLegend). All intracellular antibodies were diluted 1:200 in Perm/Wash Buffer (BD Biosciences). Cells were acquired on an LSRFortessa flow cytometer and data were analyzed with FlowJo v10. Cells were gated on live singlets prior to downstream analyses.

### Histology

Tissues were fixed in 10% neutral-buffered formalin (Fisher Scientific) and stored at 4°C until further processed. Tissue pieces were embedded in paraffin, sectioned by standard histological techniques and stained with hematoxylin and eosin. Slides were scanned on an Aperio VERSA using brightfield at 20x magnification. Histopathological disease scoring was performed by blinded board-certified pathologists. Tissue sections were given a score from 0-4 (healthy-severe) for multiple parameters, including degree of inflammatory cell infiltration, necrosis, and free bacterial colonies, along with tissue-specific parameters such as villus blunting and crypt hyperplasia.

### Statistics

Statistical analyses were performed using Prism v9.0 (GraphPad Software). Independent groups were compared by Mann-Whitney U test or Kruskal-Wallis test with Dunn’s multiple comparisons test. Survival curves were compared by Mantel-Cox test. Statistical significance is denoted as * (p<0.05), ** (p<0.01), *** (p<0.001), **** (p<0.0001), or ns (not significant).

## Acknowledgements

We thank Enrico Radaelli for constructive input and the staff at the PennVet Comparative Pathology Core for their help in preparing and analyzing the histological samples. We thank members of the Brodsky and Shin labs for scientific discussion. This work was supported by NIH Awards R01AI128530 (IEB), R01AI1139102A1 (IEB), and R01DK123528 (IEB); BWF Investigator in the Pathogenesis of Infectious Disease Award (IEB, SS); Mark Foundation Grant 19-011MIA (IEB), the Foundation Blanceflor Postdoctoral Scholarship (DS), the Swedish Society for Medical Research postdoctoral fellowship (DS) and the Sweden-America Foundation J. Sigfrid Edström award (DS); NIH NRSA F31AI160741-01 (RM); NIH T32 AI141393-2 (RM); F32 AI164655 (JPG); NIH NRSA F31AI161319 (BH); and NSF GRFP Award (SP); NIH T32 AI141393-03 (JZ); NIH Awards R21AI151476 (SS), R01AI118861 (SS), and R01AI123243 (SS). The veterinary pathologists performing the histopathological analysis are supported by the Abramson Cancer Center Support Grant (P30 CA016520). The scanner used for whole slide imaging and the image visualization software was supported by an NIH Shared Instrumentation Grant (S10 OD023465-01A1). Model figure was created using Biorender.

---

**Supplemental Figure 1. Effects of TNFR1-deficiency on pyogranuloma formation in intestine and lymphatic tissue during *Yersinia* infection**

(A) Total number of intestinal lesions at day 5 post-infection with *Yp*. Each circle represents one mouse. Lines represent median. Pooled data from four independent experiments.

(B) Flow cytometry plots displaying the gating strategy employed to identify neutrophils (CD11b+ Ly-6G+), monocytes (CD64^+^ Ly-6C^hi^), and macrophages (CD64^+^ Ly-6C^lo^ MHC-II^hi^) in small intestinal PG+ tissue. Representative images of three independent experiments.

(C) Frequencies of indicated cell types in blood of uninfected chimeric mice. Pooled data from two independent experiments.

All statistical analyses by Mann-Whitney U test. *p<0.05, **p<0.01, ***p<0.001, ****p<0.0001, ns = not significant.

**Supplemental Figure 2. Autocrine TNF signaling in monocytes is required for systemic control of *Yersinia***

(A) Frequency of indicated cell types in the blood of uninfected chimeric mice.

(B) Bacterial burdens in indicated organs at day 5 post-infection. Each circle represents one mouse. Lines represent geometric mean.

(C) Bacterial burdens in small-intestinal PG-and PG+ tissue at day 5 post *Yp*-infection. Each symbol represents one mouse. Lines represent geometric mean.

All data pooled from two independent experiments. Statistical analysis by Kruskal-Wallis test with Dunn’s multiple comparisons correction. Mann-Whitney U test. *p<0.05, **p<0.01, ***p<0.001, ****p<0.0001, ns = not significant.

**Supplemental Figure 3. TNFR1 signalling in monocytes is independent of YopJ-induced RIPK1 kinase activity**

(A) Frequency of indicated cell types in the blood of uninfected chimeric mice.

(B) Survival of *wild-type* (left) and *Tnfr1^-/-^* (right) mice infected with WT (white circles), *ΔyopE* (blue) or YopH^R409A^ (white squares) *Yp*. n = 5-32 (*wild-type*) and 13-20 (*Tnfr1^-/-^*) mice per group. Pooled data from 2-4 independent experiments.

Statistical analysis by Mantel-Cox test (B). *p<0.05, **p<0.01, ***p<0.001, ****p<0.0001, ns = not significant.

**Supplemental Figure 4. Cytokine production downstream of TNFR1 expression on monocytes is specific to IL-1 in intestinal pyogranulomas**

(A) Cytokine levels in homogenates of tissue punch biopsies were measured by cytometric bead array at day 5 post-infection of chimeric WT mice reconstituted with indicated cells. Lines represent median. Pooled data from two independent experiments.

(B) Cytokine levels in serum were measured by cytometric bead array at day 5 post-infection of chimeric WT mice reconstituted with indicated cells. Lines represent median. ND = not detected. Pooled data from two independent experiments.

(C) Frequencies of indicated cell types in small intestinal PG+ tissue or spleen at day 5 post-infection of WT chimeric mice reconstituted with the indicated cells. Pooled data from two independent experiments.

(D) Flow cytometry plots of intracellular IL-1 in monocytes (CD64^+^ Ly-6C^hi^) from small intestinal PG+ tissue in WT and *Il1b^-/-^*mice at day 5 post-infection. Plots representative of two independent experiments.

(E) Intracellular levels of TNF in monocytes and neutrophils in small intestinal PG+ tissue at day 5 post-infection. Each circle represents the mean of 3-10 pooled punch biopsies from one mouse. Lines connect congenic cell populations within individual mice. Pooled data from two independent experiments.

Statistical analyses by Kruskal-Wallis test with Dunn’s multiple comparisons correction (A, B), or congenic cells within mice: Wilcoxon test; across groups: Mann-Whitney U test (E). ns = not significant.

**Supplemental Figure 5. Systemic bacterial burdens are comparable in WT and IL-1-deficient mice.**

Bacterial burdens in indicated organ at day 5 post-infection. Each circle represents one mouse. Lines represent geometric mean. Pooled data from three independent experiments.

All statistical analyses by Mann-Whitney U test. *p<0.05, ns = not significant.

**Supplemental Figure 6. Monocyte-derived IL-1 signals to non-hematopoietic cells to restrict *Yersinia* infection**

(A) Frequencies of cell types in the blood of chimeric mice. Pooled data from two independent experiments

(B) Bacterial burdens in indicated organ at day 5 post-infection of indicated chimeric mouse. Each circle represents one mouse. Lines represent geometric mean. Pooled data from two independent experiments.

(C). Frequencies of indicated cell types in small intestinal PG+ tissue at day 5 post-infection of indicated chimeric mouse. Pooled data from three independent experiments Statistical analysis by Kruskal-Wallis test with Dunn’s multiple comparisons correction *p<0.05, ns = not significant.

(D).

## Abbreviations

CCR2: CC chemokine receptor 2
MLN: Mesenteric lymph nodes
PG: Pyogranuloma
PP: Peyer’s patches
RIPK1: Receptor-interacting protein kinase 1
TNF: Tumor necrosis factor
TNFR1: Tumor necrosis factor receptor 1
*Yp*: Yersinia pseudotuberculosis
Yop: Yersinia outer protein

