## Supplemental Figures for "A TNF-IL-1 circuit controls *Yersinia* within intestinal granulomas"

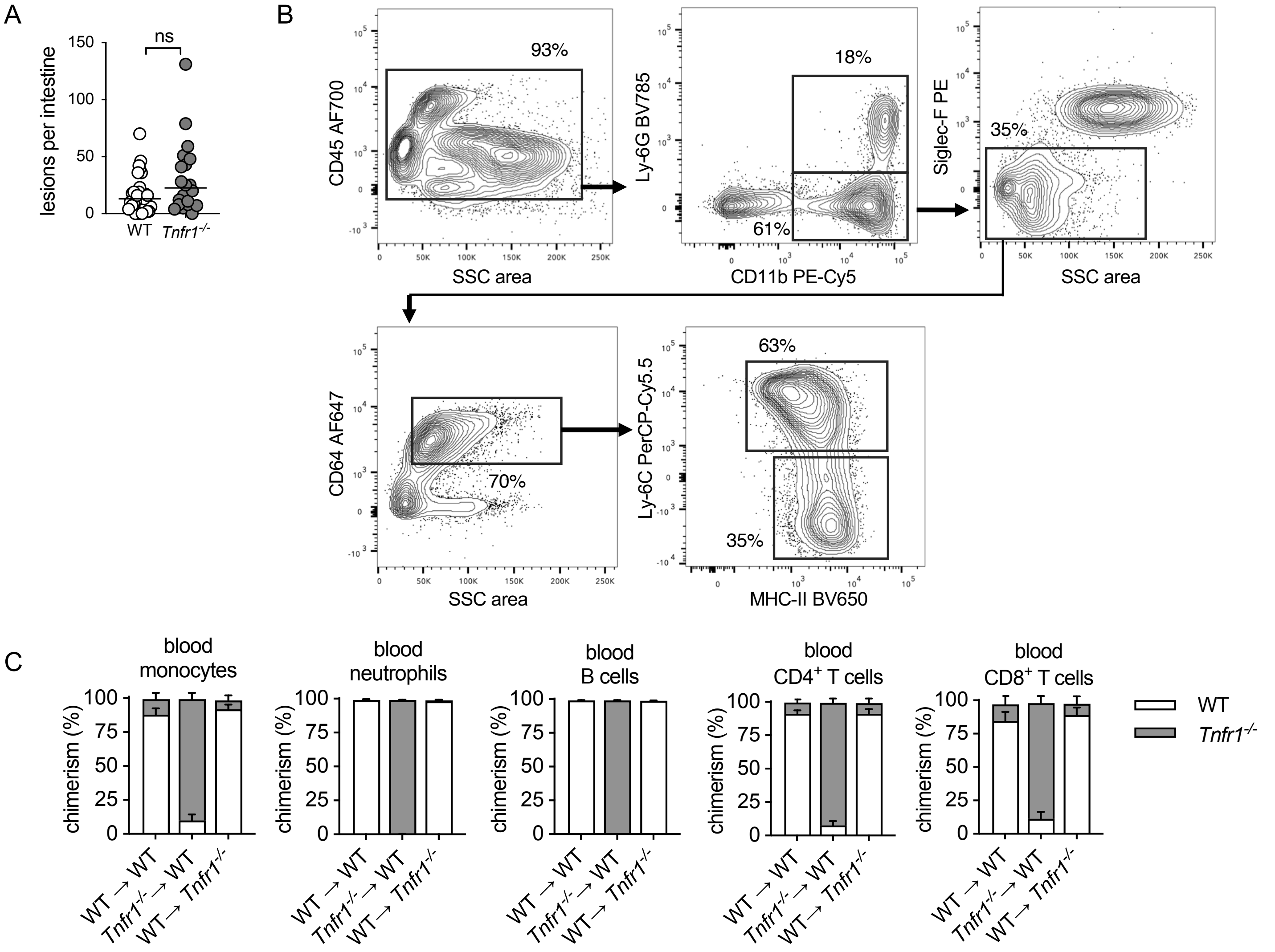


**Supplemental Figure 1. Effects of TNFR1-deficiency on pyogranuloma formation in intestine and lymphatic tissue during *Yersinia* infection**

(**A**) Total number of intestinal lesions at day 5 post-infection with *Yp*. Each circle represents one mouse. Lines represent median. Pooled data from four independent experiments.

(**B**) Flow cytometry plots displaying the gating strategy employed to identify neutrophils (CD11b+ Ly-6G+), monocytes ($CD64^{+}$ $Ly\text{-}6C^{hi}$), and macrophages ($CD64^{+}$ $Ly\text{-}6C^{lo}$ $MHC\text{-}II^{hi}$) in small intestinal PG+ tissue. Representative images of three independent experiments.

(**C**) Frequencies of indicated cell types in blood of uninfected chimeric mice. Pooled data from two independent experiments.

All statistical analyses by Mann-Whitney U test. *p<0.05, **p<0.01, ***p<0.001, ****p<0.0001, ns = not significant.

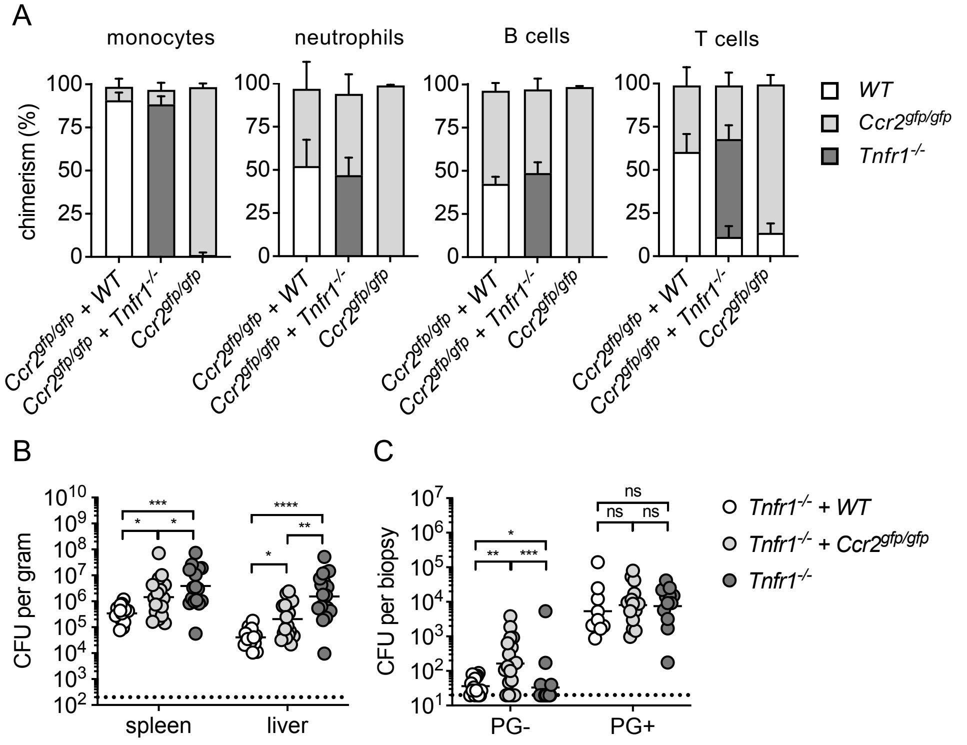


**Supplemental Figure 2. Autocrine TNF signaling in monocytes is required for systemic control of *Yersinia***
(**A**) Frequency of indicated cell types in the blood of uninfected chimeric mice.
(**B**) Bacterial burdens in indicated organs at day 5 post-infection. Each circle represents one mouse. Lines represent geometric mean.
(**C**) Bacterial burdens in small-intestinal PG- and PG+ tissue at day 5 post *Yp*-infection. Each symbol represents one mouse. Lines represent geometric mean.
All data pooled from two independent experiments. Statistical analysis by Kruskal-Wallis test with Dunn's multiple comparisons correction. Mann-Whitney U test. *p<0.05, **p<0.01, ***p<0.001, ****p<0.0001, ns = not significant.

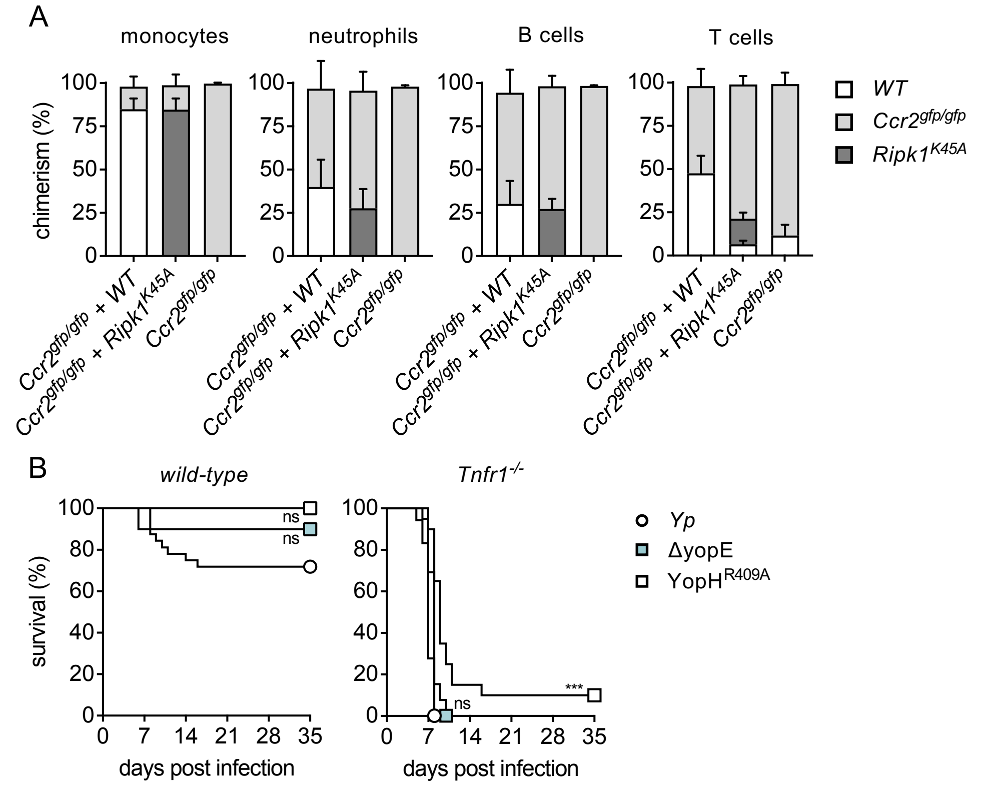


**Supplemental Figure 3. TNFR1 signalling in monocytes is independent of YopJ-induced RIPK1 kinase activity**

(**A**) Frequency of indicated cell types in the blood of uninfected chimeric mice.

(**B**) Survival of *wild-type* (left) and *Tnfr1*$^{-/-}$ (right) mice infected with WT (white circles), *ΔyopE* (blue) or YopH$^{R409A}$ (white squares) *Yp*. n = 5-32 (*wild-type*) and 13-20 (*Tnfr1*$^{-/-}$) mice per group. Pooled data from 2-4 independent experiments.

Statistical analysis by (B) Mantel-Cox test. *p<0.05, **p<0.01, ***p<0.001, ****p<0.0001, ns = not significant.

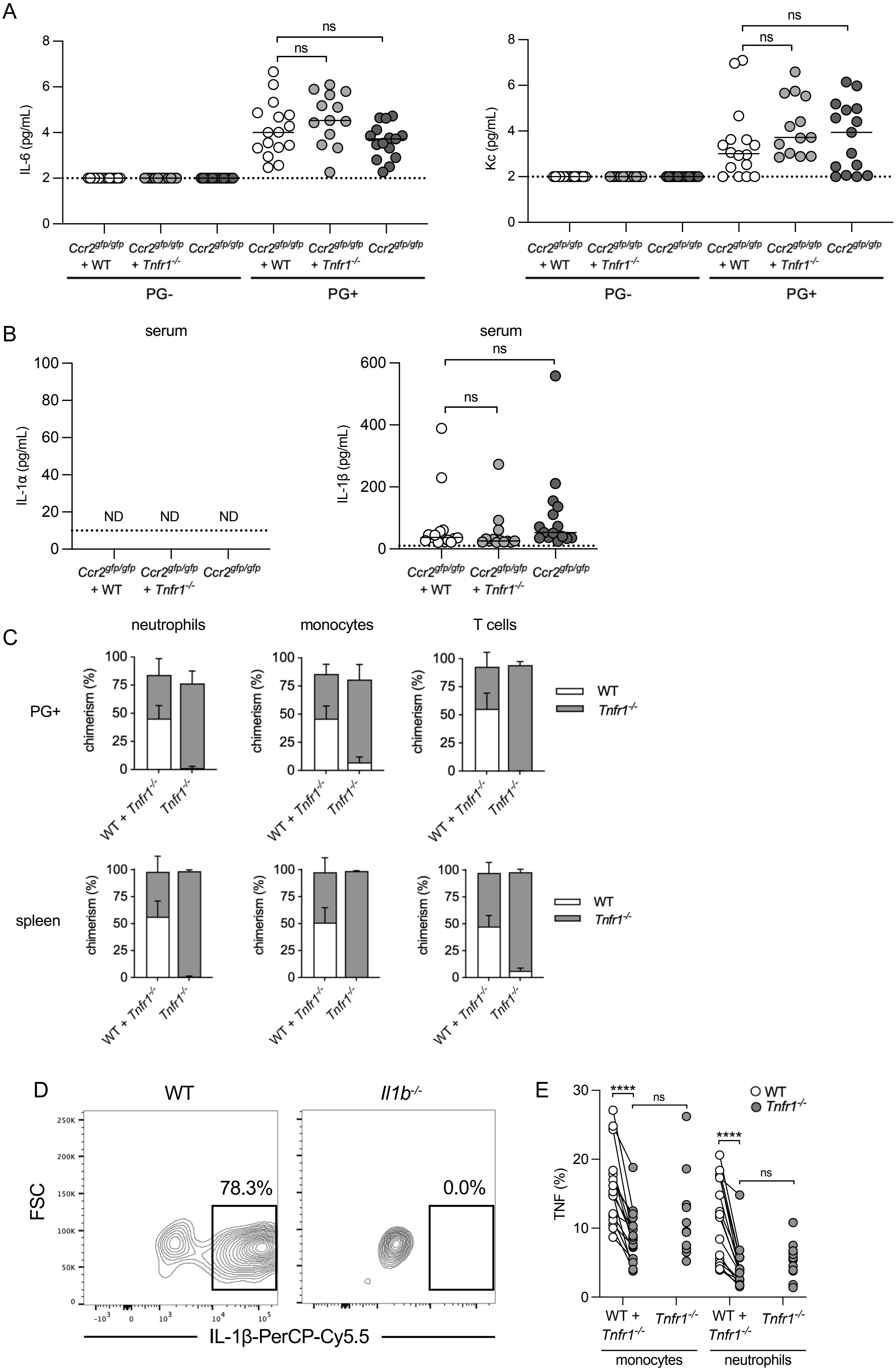
A
IL-6 (pg/mL)
Kc (pg/mL)
ns
Ccr2gfp/gfp + WT
Ccr2gfp/gfp + Tnfr1-/-
Ccr2gfp/gfp
PG-
PG+
B
serum
IL-1α (pg/mL)
ND
IL-1β (pg/mL)
C
neutrophils
monocytes
T cells
PG+
spleen
chimerism (%)
WT + Tnfr1-/-
Tnfr1-/-
WT
D
WT
Il1b-/-
FSC
78.3%
0.0%
IL-1β-PerCP-Cy5.5
E
TNF (%)
****
monocytes
neutrophils

**Supplemental Figure 4. Cytokine production downstream of TNFR1 expression on monocytes is specific to IL-1 in intestinal pyogranulomas**

(**A**) Cytokine levels in homogenates of tissue punch biopsies were measured by cytometric bead array at day 5 post-infection of chimeric WT mice reconstituted with indicated cells. Lines represent median. Pooled data from two independent experiments.

(**B**) Cytokine levels in serum were measured by cytometric bead array at day 5 post-infection of chimeric WT mice reconstituted with indicated cells. Lines represent median. ND = not detected. Pooled data from two independent experiments.

(**C**) Frequencies of indicated cell types in small intestinal PG+ tissue or spleen at day 5 post-infection of WT chimeric mice reconstituted with the indicated cells. Pooled data from two independent experiments.

(**D**) Flow cytometry plots of intracellular IL-1 in monocytes ($CD64^{+}$ Ly-$6C^{hi}$) from small intestinal PG+ tissue in WT and *Il1b*$^{-/-}$ mice at day 5 post-infection. Plots representative of two independent experiments.

(**E**) Intracellular levels of TNF in monocytes and neutrophils in small intestinal PG+ tissue at day 5 post-infection. Each circle represents the mean of 3-10 pooled punch biopsies from one mouse. Lines connect congenic cell populations within individual mice. Pooled data from two independent experiments.

Statistical analyses by Kruskal-Wallis test with Dunn’s multiple comparisons correction (A, B), or congenic cells within mice: Wilcoxon test; across groups: Mann-Whitney U test (E). ns = not significant.

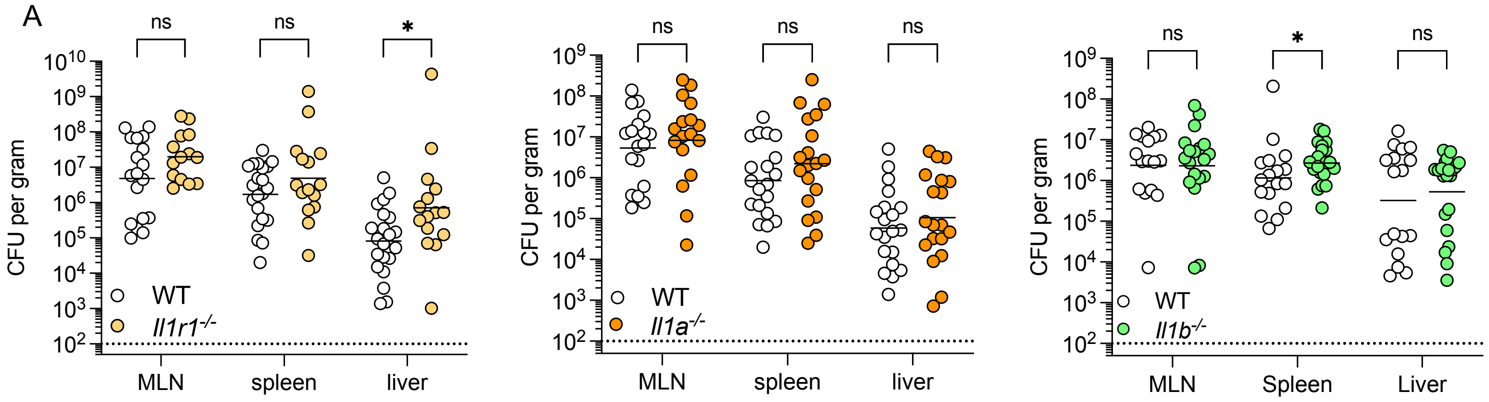


**Supplemental Figure 5. Systemic bacterial burdens are comparable in WT and IL-1-deficient mice.**
Bacterial burdens in indicated organ at day 5 post-infection. Each circle represents one mouse. Lines represent geometric mean. Pooled data from three independent experiments.
All statistical analyses by Mann-Whitney U test. *p<0.05, ns = not significant.

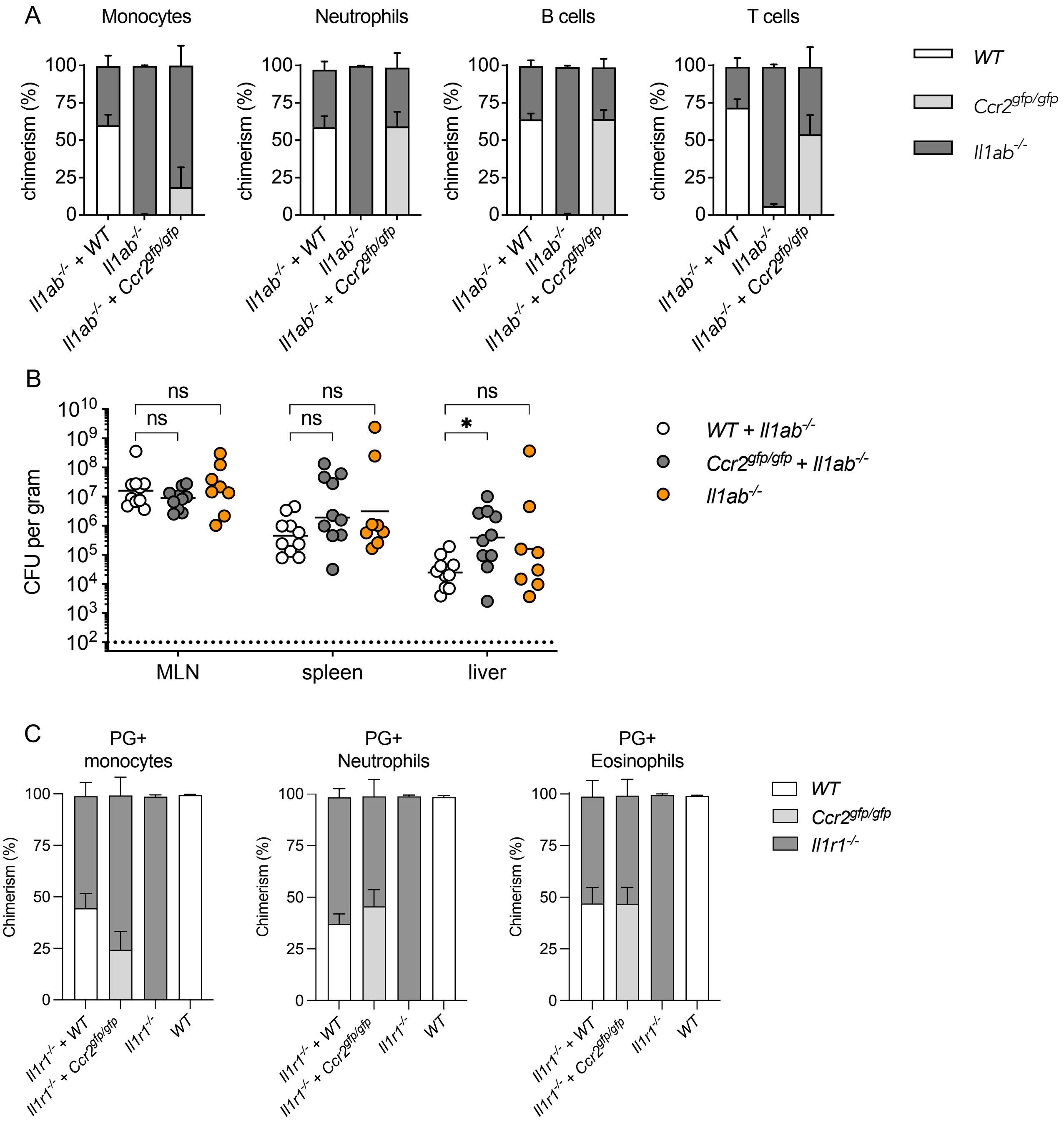


**Supplemental Figure 6. Monocyte-derived IL-1 signals to non-hematopoietic cells to restrict *Yersinia* infection**

(A) Frequencies of cell types in the blood of chimeric mice. Pooled data from two independent experiments

(B) Bacterial burdens in indicated organ at day 5 post-infection of indicated chimeric mouse. Each circle represents one mouse. Lines represent geometric mean. Pooled data from two independent experiments.

(C) Frequencies of indicated cell types in small intestinal PG+ tissue at day 5 post-infection of indicated chimeric mouse. Pooled data from three independent experiments

Statistical analysis by (B) Kruskal-Wallis test with Dunn's multiple comparisons correction. *p<0.05, ns = not significant.
